# Group comparison based on genetic information reveals lineage-specific therapeutic vulnerabilities in acute myeloid leukemia

**DOI:** 10.1101/2023.05.18.541265

**Authors:** Jakushin Nakahara, Keita Yamamoto, Tomohiro Yabushita, Takumi Chinen, Kei Ito, Yutaka Takeda, Daiju Kitagawa, Susumu Goyama

## Abstract

Cancer is a genetic disease with specific mutations or fusions. Therapies targeting cancer cell-specific essential genes are expected to have efficient anticancer effects with fewer side effects. To explore such cancer cell-specific vulnerabilities, we established a two-group comparison system to predict essential genes in each cancer subtype using the data from the Cancer Dependency Map (DepMap). We applied this analytical method to acute myeloid leukemia (AML) and identified PCYT1A and BCL2L1 as a specific vulnerability in MLL-rearranged AML and *TP53*-mutated AML, respectively. Interestingly, further investigation revealed that PCYT1A is in fact a critical regulator in monocytic AML including those with MLL-rearrangements, and BCL2L1 is essential in acute erythroid leukemia in which *TP53* is frequently mutated. These results highlighted the importance of cell of origin, rather than the genetic aberrations alone, to identify subtype-specific vulnerabilities in AML. The DepMap-based two-group comparison approach could accelerate the discovery of subtype-specific therapeutic targets in diverse cancers.

## INTRODUCTION

Cancer is a genetic disease caused by changes in genes involved in cell division, cell death, proliferation and differentiation^1, 2^. The genetic alterations in cancer cells lead to distinct gene expression, which provides cancer-specific vulnerability genes^3^. Compared to conventional chemotherapy, molecular targeted therapies targeting such cancer-specific vulnerability generally have efficient anticancer effects with fewer side effects^4, 5^. Over the past two decades, the number of targeted drugs available for cancer therapy has steadily increased. However, due to the diversity of genetic abnormalities in cancer cells, there remains an unmet medical need to explore therapeutic targets for each type of cancer.

The Cancer Dependency Map (DepMap) is an ongoing project to identify essential genes in over 1000 cancer cell lines through genome-wide CRISPR and shRNA screening^6, 7^. DepMap provides a unique Gene Effect Score (also called Chronos Score in the latest version), which represents the essentiality of a gene in each cancer cell. The lower the Gene Effect Score, the more essential the gene is for the survival and proliferation of the cell, with a value of 0 indicating that the gene is not essential and a value of -1 indicating that the gene is essential. By comparing the Gene Effect Score for each cancer type, we can extract genes that are specifically essential for a particular cancer type^8^. In fact, numerous recent studies have identified potential therapeutic targets in specific cancer types using the DepMap information^9-12^. However, genetic alterations vary among patients and cell lines even within a specific cancer type, and it is not possible to extract essential genes according to the genetic classification on the DepMap portal website^6^. We therefore developed a system to extract essential genes more efficiently by grouping mutated genes or fusion genes within a specific cancer cell type, which enables us to extract specific therapeutic vulnerabilities in each cancer subtype with a specific mutation or a fusion.

Acute myeloid leukemia (AML) is a blood cancer with unique chromosomal translocations and gene mutations^13, 14^. The genetic heterogeneity in AML is one of the main reasons for failure of classic chemotherapy. Certain genetic aberrations have been shown to be associated with inferior outcomes of AML. One of these is a balanced rearrangement involving the *KMT2A* (also called MLL) gene, located at 11q23. *MLL* fuses with more than 100 partner genes, including MLLT3 (AF9) and MLLT1 (ENL), creating fusion proteins with leukemogenic activity^15^. The MLL-rearranged AML is often associated with poor prognosis, monocytic or myelomonocytic differentiation, and constitutive activation of MLL target genes, such as HOXA9 and MEIS1^16-20^. Another genetic aberration associated with poor outcome of AML are mutations in *TP53* gene^21-23^. *TP53* is a critical tumor suppressor gene and are frequently mutated in AML related to an increased genomic instability, such as therapy‐related AML and AML with myelodysplasia. *TP53* mutations are also frequently detected in acute erythroid leukemia (AEL), a rare and aggressive subtype of AML^24, 25^.

In this study, we applied our two-group comparison approach to identify specific therapeutic targets in AMLs with MLL-rearrangements or *TP53* mutations. The *in-silico* analyses and subsequent investigation using AML cell lines led to the identification of PCYT1A and BCL2L1 as a critical regulator in monocytic AML and AEL, respectively.

## METHODS

### The DepMap-based two-group comparison system

The system is written in Python on Google Colaboratory and performs automatic and fast analysis and display of results by entering parameters in the Python code. The analysis flow is as follows. First, download the following files from the DepMap Public 22Q2 database (DepMap Public dataset which was released on May 22nd, 2022): cell classification information, Gene Effect Score, gene expression levels, gene mutations, and gene fusions. We can then select whether we want to compare Gene Effect Score or expression levels, and further select two groups of cell lines. There are six ways to select cell groups: primary disease, lineage, lineage-subtype, lineage-sub-subtype, all cells, and manual entry. After two cell line groups have been selected in this way, we can choose whether to further narrow this group of cell lines by gene mutation or fusion by entering the name of the gene. The cell lines in the group will then be limited to those with the gene mutation or fusion. The two groups of cell lines to be compared are now defined. The data are then automatically and quickly analyzed by Python and the results are displayed. This analysis first calculates the average Gene Effect Score or expression level of each gene within each cell line group. Then, the difference of the mean values of each gene between the two groups is calculated, and the genes are sorted and ranked in the order of the difference (Fig. 1a). At the same time, unpaired t test with Welch’s correction and Benjamini-Hochberg procedure are performed for each gene between the two groups, and the p values and q values are calculated. Through these processes, genes with large differences in Gene Effect Score or expression levels between the two groups of cell lines are extracted. Genes with particularly low Gene Effect Score in one group may be a specific drug target in that cancer. Therefore, this system can extract target candidates that are specific to a group of cell lines with a particular gene mutation or fusion. The system can be used by downloading the code from GitHub repository and running it on Google Colaboratory (Nakahara, 2023, *GitHub*. Available at: https://github.com/jakushinn/depmap_analysis)^26^.

**Fig. 1.**
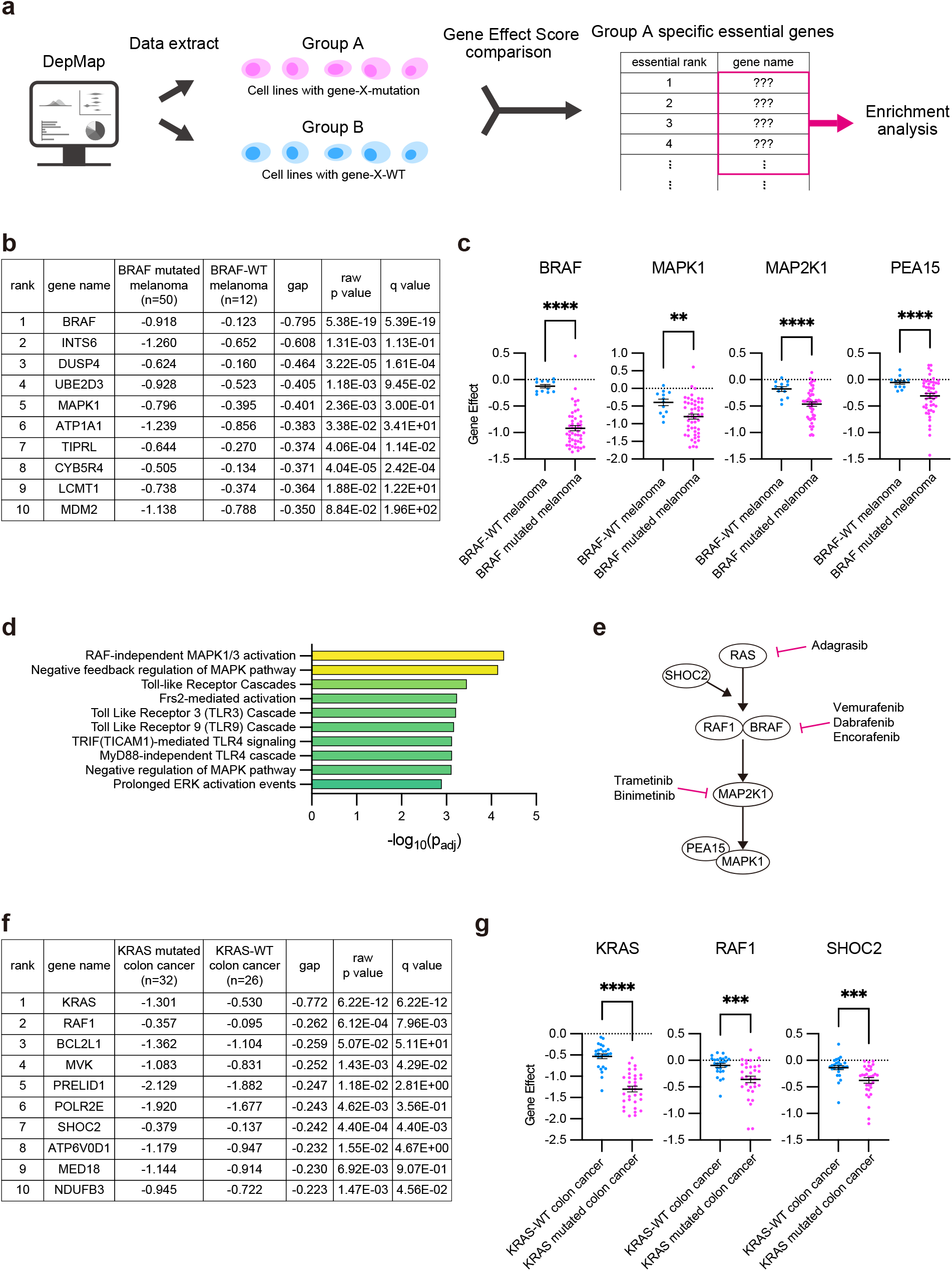
DepMap Public dataset-based two-group comparison to identify specific essential genes. (**a**) Schematic presentation of the two-group comparison system using the DepMap information. Two groups of cell lines are defined by cancer types and mutated genes or fusion genes. The averages of each Gene Effect Score are compared between two groups, and the genes are sorted by the rank of the group-A-specific essentiality. Enrichment analysis of the top-ranked genes can reveal the essential pathways or cascades in group A. (**b**) Results of DepMap Gene Effect Score analysis between *BRAF*-mutated and *BRAF*-WT melanoma (*BRAF*-mutated melanoma: n=50, *BRAF*-WT melanoma: n=12). The top10 genes that are specifically essential in *BRAF*-mutated melanoma are shown in the table. (**c**) Dot plots of Gene Effect Scores of BRAF, MAPK1, MAP2K1, and PEA15 for *BRAF*-mutated or WT melanoma. (**d**) Reactome enrichment analysis for the top 30 essential genes in *BRAF*-mutated melanoma using g:Profiler (https://biit.cs.ut.ee/gprofiler). (**e**) Schematic presentation of the MAPK pathway. The genes shown are specifically essential in *BRAF*-mutated melanoma or *KRAS*-mutated colon cancer (**c, g**). (**f**) Results of DepMap Gene Effect Score analysis between *KRAS*-mutated and *KRAS*-WT colon cancer. (*KRAS*-mutated colon cancer: n=32, *KRAS*-WT colon cancer: n=26) The top10 genes which are specifically essential in *KRAS*-mutated colon cancer are shown in the table. (**g**) Dot plots of Gene Effect Scores of KRAS, RAF1, and SHOC3 for colon cancer. Data are shown as mean ± s.e.m. \*\**P*<0.01, \*\*\**P*<0.001, \*\*\*\**P*<0.0001, unpaired t test with Welch’s correction and Benjamini-Hochberg procedure.

### Plasmids and viral infection

Annealed oligonucleotides were cloned into the pLKO5.sgRNA.EFS.tRFP657 vector, pLKO5.sgRNA.EFS.GFP vector or lentiGuide-Puro vector to generate single-guide (sg)RNA expression vector. pLKO5.sgRNA.EFS.tRFP657 (Addgene plasmid # 57824 ; http://n2t.net/addgene:57824 ; RRID:Addgene_57824) and pLKO5.sgRNA.EFS.GFP (Addgene plasmid # 57822 ; http://n2t.net/addgene:57822 ; RRID:Addgene_57822) were gifts from Benjamin Ebert.^27^ lentiGuide-Puro was a gift from Feng Zhang (Addgene plasmid # 52963 ; http://n2t.net/addgene:52963 ; RRID:Addgene_52963).^28^ The Cas9 expression in THP1, MOLM13, HEL and KASUMI1 was induced by lentiCas9-Blast, and the Cas9 expression in TF1 and OCI-AML3 was induced by FUCas9Cherry. lentiCas9-Blast was a gift from Feng Zhang (Addgene plasmid # 52962 ; http://n2t.net/addgene:52962 ; RRID:Addgene_52962).^28^ FUCas9Cherry was a gift from Marco Herold (Addgene plasmid # 70182 ; http://n2t.net/addgene:70182 ; RRID:Addgene_70182).^29^ Lentiviruses were produced by transient transfection of 293T cells with viral plasmids along with gag-, pol-, and env-expressing plasmids (pMD2.G and psPAX) using the calcium-phosphate method.^30^ pMD2.G (Addgene plasmid #12259; http://n2t.net/addgene:12259; RRID: Addgene_12259) and psPAX2 (Addgene plasmid #12260; http://n2t.net/addgene:12260; RRID: Addgene_12260) were gifts from Didier Trono. Sequence of the nontargeting (NT) control and sgRNAs targeting for human PCYT1A, mouse Pcyt1a, human PCYT1B, human TP53 and human BCL2L1 from 5′ to 3′ are provided as follows: CGCTTCCGCGGCCCGTTCAA (NT), ATTATGATGTGTATGCGAGG (sgPCYT1A#1), AATACGTACCTCATTGTGGG (sgPCYT1A#2), CTTTAGTAAGCCCTATGTCA (sgPcyt1a#1), ACTATGATGTGTATGCAAGA (sgPcyt1a#2), GAGCGCTGCTCAGATAGCGA (sgTP53), CAGGCGACGAGTTTGAACTG (sgBCL2L1#1), GACCCCAGTTTACCCCATCC (sgBCL2L1#2).

### The engineered human and mouse AML cells

Human and mouse MLL-AF9-expressing AML cells were generated in previous studies by transducing MLL-AF9 into human cord blood CD34^+^ cells and mouse bone marrow progenitor cells, respectively, using retroviruses^31^. The cSAM cells were generated in previous studies by transducing mutant ASXL1 (ASXL1-E635RfsX15) and mutant SETBP1 (SETBP1-D868N) into mouse bone marrow progenitor cells using retrovirus ^31-34^ All animal studies were approved by the Animal Care Committee of the Institute of Medical Science at the University of Tokyo (approval number: PA22-17, PA22-23), and were conducted in accordance with the Regulation on Animal Experimentation at University of Tokyo based on International Guiding Principles for Biomedical Research Involving Animals.

### Cell culture

THP1, MOLM13, KASUMI1, HEL, OCIAML3, and U937 cells were cultured in RPMI-1640 with 10% fetal bovine serum (FBS). TF1 cells were cultured in RPMI-1640 with 10% FBS and 1 ng/mL recombinant human granulocyte–macrophage colony-stimulating factor (rhGM-CSF, No. 215-GM, R&D Systems, Minneapolis, MN). 293T cells were cultured in DMEM media containing 10% FBS. The mouse MLL-AF9-transduced cells were cultured in MethoCult™ M3234 (STEMCELL Technologies, Bancouber, BC, Canada). The human MLL-AF9-transduced cells were cultured in StemSpan SFEM II medium (#ST-09655, STEMCELL Technologies) containing 1% penicillin– streptomycin together with 10 ng/ml rhSCF (#255-SC, R&D Systems), 10 ng/ml rhTPO (#288-TP, R&D Systems), 10 ng/ml rmFlt-3 ligand (#427-FL, R&D Systems), 10 ng/ml rhIL-3 (#203-IL, R&D Systems) and 10 ng/ml rhIL-6 (#206-IL, R&D Systems).^30^

### Cell viability assay

Cells were seeded in a 96-well plate at a density of 1 × 10^4^ cells per well and were cultured with A-1155463 for 72 h.^35^ Cell viability was measured by Cell Counting Kit-8 (Dojindo, Kumamoto, Japan) according to the manufacturer’s protocols. The absorbance was measured with CLARIOstar PLUS (BMG LABTECH, Ortenberg, Germany).

### Western blot analysis

Cells were lysed in sample buffer (2× Laemmli sample buffer, No. 1610737, Bio-Rad, Hercules, California, USA). Whole-cell lysates were subjected to sodium dodecyl sulfate polyacrylamide gel electrophoresis (SDS-PAGE) and transferred to a polyvinylidene fluoride membrane (Bio-Rad). The blots were incubated with anti-CCTα (D18B6) antibody (#6931; Cell Signaling Technology, Beverly, Massachusetts, USA), anti-PCYT1B antibody (#13765-1-AP; Thermo Fisher Scientific, Waltham, Massachusetts, USA), anti-p53 antibody (#sc-126; Santa Cruz Biotechnology, Dallas, Texas, USA), anti-Bcl-xL(H-5) antibody (#sc-8392; Santa Cruz Biotechnology, Dallas, Texas, USA) and anti-GAPDH antibody (#5174; Cell Signaling Technology, Beverly, Massachusetts, USA). Signals were detected with ECL Western Blotting Substrate (Promega Corp., Fitchburg, Wisconsin, USA), and immunoreactive bands were visualized using ChemiDoc Touch MP (Bio-Rad, Hercules, California, USA).

### Flow cytometry

Expression of mCherry, tRFP657 and GFP was analyzed with a FACS CytoFLEX (Beckman Coulter, Brea, California, USA). The data were analyzed using FlowJo software (Treestar, Inc., San Carlos, California, USA).

### Statistical analyses

GraphPad Prism 9 was used for statistical analyses. Unpaired Welch’s *t*-test (two-tailed) and Ordinary two-way ANOVA were used for pairwise comparisons of significance. A *P*-value > 0.05 was considered as not significant (ns). Sample size was decided based on our previous experience in the field, not predetermined by a statistical method.

## RESULTS

### Development of a DepMap-based two-group comparison system to identify therapeutic vulnerabilities in specific cancer subtypes

We first developed a method to group cancer cells based on the information in DepMap, including cell classification, gene mutations, and gene fusions. In this system, we can define the two groups of cell lines to be compared based on the type of cancer (e.g., primary disease or lineage-subtype) and genetic aberration (gene mutation or gene fusion). The Gene Effect Score or gene expression levels in each group are automatically and quickly analyzed by Python to calculate the difference of the mean values of each gene between the two groups. The genes are then ranked in the order of the difference with statistical analysis (Fig. 1a). Genes with particularly low Gene Effect Score in one group could be a specific drug target in a specific cancer subtype. This two-group comparison system can be used by downloading the code and preprocessed data from the GitHub repository (Nakahara, 2023, *GitHub*. Available at: https://github.com/jakushinn/depmap_analysis) and running it on a local computer.

To examine the usefulness of our system, we first analyzed melanoma with/without *BRAF* mutations. We divided melanoma cells into BRAF-wild-type (WT) or BRAF-mutant cells and analyzed the Gene Effect Score to extract genes that are more demanding for cell proliferation in *BRAF*-mutated melanoma. As expected, genes in the RAS/MAPK signaling pathway, BRAF, MAPK1 (ERK2), MAP2K1 (MEK1), and PEA15, were ranked within the top 30 (gap of Gene Effect Score for each gene: -0.795, -0.401, -0.287, -0.254; two-tailed p value for each gene: <0.0001, 0.0024, <0.0001, <0.0001) (Fig. 1b-c, e). Reactome enrichment analysis using g:Profiler (https://biit.cs.ut.ee/gprofiler) on these top 30 ranked genes revealed critical role of RAF-independent MAPK1/3 activation (Fig. 1d)^36, 37^. These results indicate that *BRAF*-mutated melanoma is dependent on the RAS/MAPK signaling pathway^38, 39^. In fact, BRAF (vemurafenib, dabrafenib, and encorafenib) and MAP2K1 (trametinib and binimetinib) inhibitors have been used to treat *BRAF*-mutated melanoma in the clinical setting (Fig. 1e)^40-44^.

We next applied our system to *KRAS*-WT/mutated colon cancers to identify the therapeutic targets in colon cancers with *KRAS* mutations. Genes in the RAS pathway ranked high, and especially, KRAS itself ranked first (gap of Gene Effect Score, -0.772; two-tailed p value, <0.0001) (Fig. 1e-g). In the clinical setting, the KRAS inhibitor adagrasib (MRTX849) has been reported to be effective on *KRAS*-mutated colon cancers in a phase 1 trial^45, 46^. These results also indicate the usefulness of the two-group comparison approach to identify therapeutic targets that are highly specific to a cancer subtype with a specific mutation.

### Exploring therapeutic vulnerabilities in MLL-rearranged AML

Next, we used our two-group comparison method to identify novel therapeutic targets in MLL-rearranged AML. The DepMap Public 22Q2 database contains Gene Effect Scores of eight AML cell lines with MLL rearrangements and eighteen AML cell lines with wild-type (WT) MLL. We compared and analyzed the Gene Effect Scores between these two groups of cell lines and extracted genes that are more important in MLL-rearranged AML. This analysis revealed several known targets for MLL-rearranged AML, such as MEN1, MBNL1 and SPI1^47-50^ (gap of Gene Effect Score, -0.583, -0.473, -0.420; two-tailed p value, 0.0023, 0.0113, 0.0134). Interestingly, the top ranked gene in our analysis was PCYT1A (gap of Gene Effect Score, -0.806; two-tailed p value, <0.0001) (Fig. 2a-b). Although the role of PCYT1A in MLL-rearranged AML has never been reported, a previous genome-wide CRISPR-Cas9 library screen using mouse MLL-AF9-expressing AML cells reported Pcyt1a as one of the dropout hits^51^. PCYT1A is a rate-limiting enzyme that synthesizes CDP-choline from phosphocholine in the phosphatidylcholine biosynthetic pathway (Kennedy pathway) (Fig. 2c)^52-55^. Other genes in the Kennedy pathway (CHKA, CHKB, PCYT1B, CEPT1, and CHPT1) showed similar Gene Effect Scores between MLL-WT and MLL-rearranged AMLs (Fig. 2d).

**Fig. 2.**
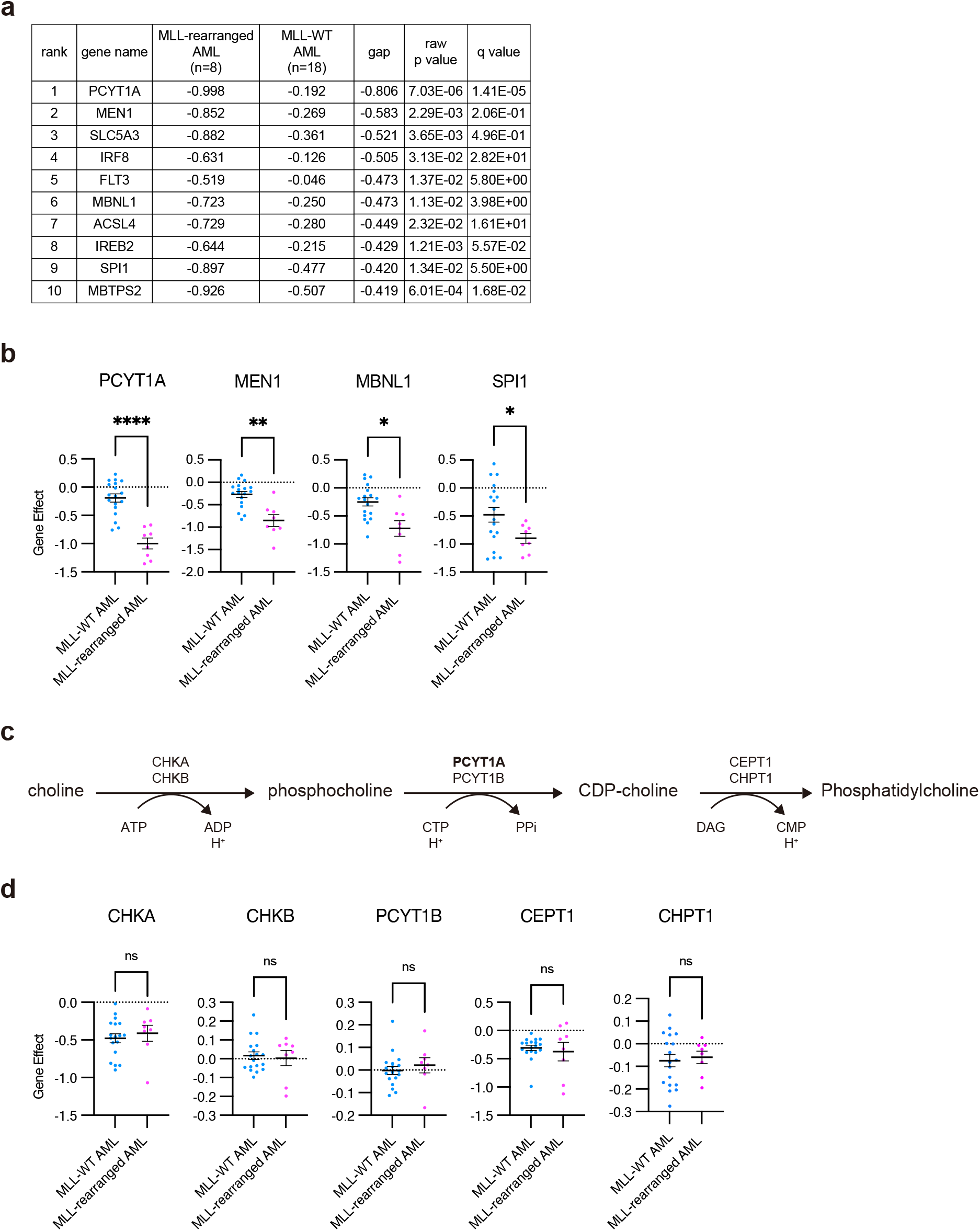
DepMap dataset analysis between MLL-rearranged and MLL-WT AML. (**a**) Results of DepMap Gene Effect Score analysis between MLL-rearranged and MLL-WT AML. (MLL-rearranged AML: n=8, MLL-WT AML: n=18) The top10 genes that are specifically essential in MLL-rearranged AML are shown in the table. (**b**) Dot plots of Gene Effect Scores of PCYT1A, MEN1, MBNL1 and SPI1 for AML. (**c**) Schematic presentation of the Kennedy pathway. (**d**) Dot plots of Gene Effect Scores of the genes involved in the Kennedy pathway for AML. Data are shown as mean ± s.e.m. ns: not significant, \**P*<0.05, \*\**P*<0.01, \*\*\*\**P*<0.0001, unpaired t test with Welch’s correction and Benjamini-Hochberg procedure.

### MLL-rearranged AML cells are dependent on PCYT1A due to the low level of PCYT1B

To confirm the importance of PCYT1A in MLL-rearranged AML, we depleted PCYT1A in two MLL-rearranged (THP1 and MOLM13) and two MLL-WT (KASUMI1 and HEL) AML cell lines using the CRISPR/Cas9 system. We transduced Cas9 [coexpressing Blasticidin S Deaminase (BSD)] together with non-targeting (NT) or single-guide (sg)RNAs targeting PCYT1A (coexpressing tRFP657) into these AML cells, and monitored the frequency of tRFP657-positive cells in culture starting 72 hours after the transduction (Fig. 3a, b)^27, 28^. Efficient depletion of PCYT1A in these cells were confirmed by western blotting (Fig. 3c). Consistent with the data in DepMap, PCYT1A depletion suppressed the growth of MLL-rearranged AML cells but not that of MLL-WT AML cells (Fig. 3d). To understand why the MLL-rearranged AML cells are dependent on PCYT1A, we assessed expression levels of six Kennedy pathway genes in AML cell lines registered in DepMap Public 22Q2. Although no genes (including PCYT1A itself) were upregulated in MLL-rearranged AML, expression of PCYT1B, a paralog of PCYT1A, was significantly lower in MLL-rearranged AML compared to MLL-WT AML (Fig. 4a). In addition, the Gene Effect Score of PCYT1A positively correlated with PCYT1B expression in AML cell lines (Pearson’s correlation coefficient, 0.6810) (Fig. 4b). These data indicate that the cells with high PCYT1B expression are less dependent on PCYT1A, while those with low PCYT1B expression are highly dependent on PCYT1A for proliferation. To examine the potential compensatory role of PCYT1B in PCYT1A-depleted AML cells, we then depleted PCYT1A, PCYT1B or both in AML cells (Fig. 4c). Depletion of PCYT1B in HEL and TF1 cells was confirmed by western blotting (Fig. 4d). Consistent with the earlier results, depletion of PCYT1A alone was sufficient to inhibit the growth of MLL-rearranged AML cell lines (THP1 and MOLM13) but not in MLL-WT AML cell lines (TF1 and HEL). Depletion of PCYT1B alone showed no effect in all the AML cell lines. Importantly, simultaneous depletion of PCYT1A and PCYT1B showed the strong growth-inhibitory effect even in TF1 and HEL cells (Fig. 4e, f), indicating that PCYT1B compensates for the loss of PCYT1A in these cells. These results suggest that MLL-rearranged AML cells are highly dependent on PCYT1A primarily because PCYT1B expression is suppressed in them.

**Fig. 3.**
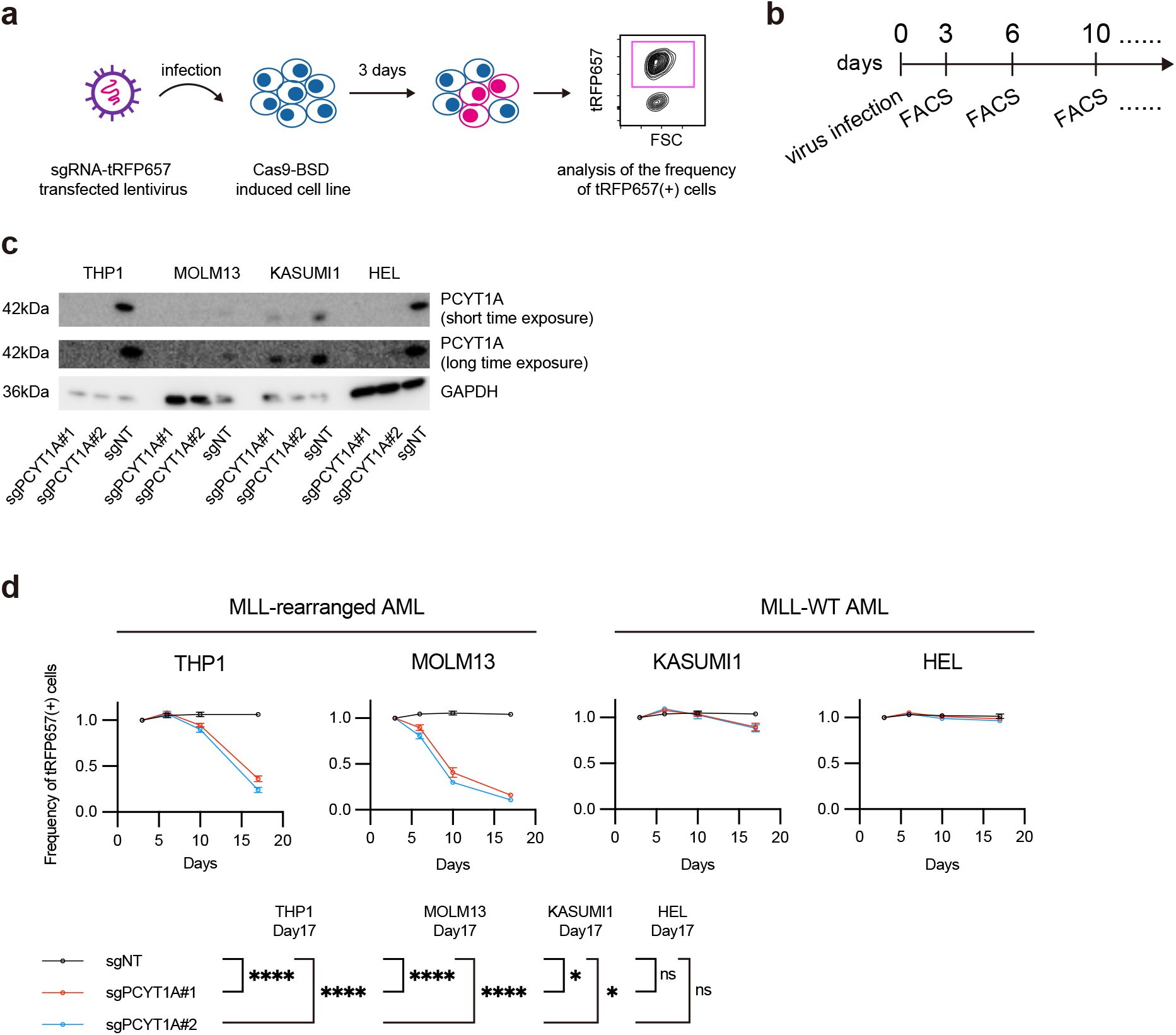
PCYT1A depletion suppresses the cell proliferation in MLL-rearranged AML. (**a-b**) Schematic presentation of experimental procedures for experiments shown in Fig. 3d. Cas9-transduced cells (coexpressing BSD) were transduced with non-targeting (NT) or single-guide (sg)RNAs targeting PCYT1A (coexpressing tRFP657) after blasticidin selection (10 μg/mL). The frequency of tRFP657-positive cells was monitored by FACS starting 72 hours after the sgRNA transduction. (**c**) PYCT1A depletion by each sgRNA was confirmed in human AML cell lines. (**d**) Changes in the frequency of tRFP657^+^ cells (sgRNA-transduced cells) in cell cultures (triplicates) are shown. Results are normalized to the frequency of tRFP657^+^ cells at day 3, set to 1. Data are shown as mean ± s.d. ns: not significant, \**P*<0.05, \*\*\*\**P*<0.0001, two-way ANOVA multiple comparisons.

**Fig. 4.**
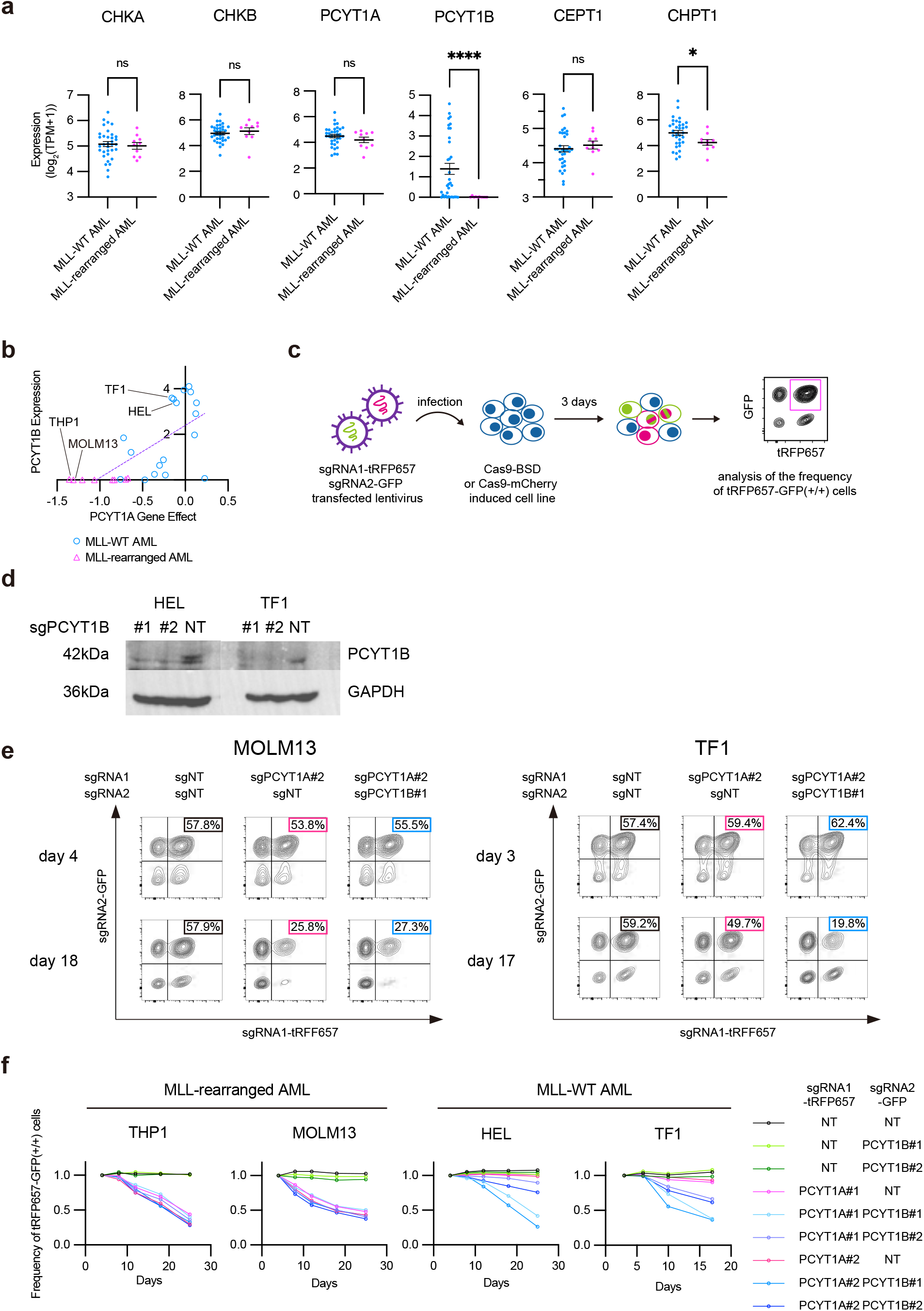
MLL-rearranged AML cells are highly dependent on PCYT1A due to the low level of PCYT1B. **(a)** Dot plots of gene expression TPM values of the Kennedy pathway genes in AML (log_2_(TPM+1)). **(b)** 2-dimentional plots of PCYT1A Gene Effect Scores and PCYT1B gene expression TPM values for AML. (n=26, pearson r: 0.6810, two-tailed p value: 0.0001) (**c**) Schematic presentation of experimental procedures for experiments shown in Fig. 4e, f. Cas9-transduced cells (coexpressing BSD) were transduced with two of NT (coexpressing tRFP657 or GFP), sgRNAs targeting PCYT1A (coexpressing tRFP657) or sgRNAs targeting PCYT1B (coexpressing GFP) after blasticidin selection (10 μg/mL). The frequency of cells that were positive for both tRFP657 and GFP was monitored by FACS starting 72 hours after the sgRNA-transduction. (**d**) PYCT1B depletion by each sgRNA was confirmed in HEL and TF1 cells. (**e**) Representative FACS plots of MOLM13 (MLL-rearranged AML) and TF1 (MLL-WT AML) at day3 and day 17 are shown. (**f**) Changes in the frequency of tRFP657^+^GFP^+^ cells (sgRNA-transduced cells) in cell cultures are shown. Results are normalized to the frequency of tRFP657^+^GFP^+^ cells at day 3, set to 1. Data are shown as mean ± s.d.

### PCYT1A is a critical regulator in monocytic AML

To further assess the role of PCYT1A in MLL-rearranged AML, we then performed similar experiments using mouse bone marrow cells and human cord blood cells transformed by a leukemogenic MLL-fusion gene, MLL-AF9^31^. We also used MLL-WT mouse AML cells transformed by ASXL1 and SETDB1 mutations (<u>c</u>ombined expression of <u>S</u>ETDB1 and <u>A</u>SXL1 <u>M</u>utations: cSAM cells^33, 34, 56^) in this experiment. As expected, depletion of PCYT1A/Pcyt1a in human or mouse MLL-AF9 cells inhibited their growth, indicating again the critical role of PCYT1A in MLL-rearranged AML. To our surprise, Pcyt1a depletion also inhibited the growth of cSAM cells despite the absence of the MLL-rearrangement. Because MLL rearrangements are common in monocytic AML [French-American-British (FAB) subtypes M4 or M5] and cSAM cells also display monocytic differentiation, we hypothesized that PCYT1A is essential for survival and proliferation in monocytic AML. Indeed, Pcyt1b level is low in cSAM cells (Fig. 5b). To test this hypothesis, we assessed the effect of PCYT1A depletion in other monocytic AML cell lines without MLL rearrangements (OCIAML3 and U937). Efficient depletion of PCYT1A in these cells was confirmed by western blotting (Fig. 5c). PCYT1A depletion inhibited the growth of these two cell lines (Fig. 5d), indicating the indispensable role of PCYT1A in monocytic AML. We then examined levels of PCYT1A and PCYT1B in primary AML cells collected from AML patients^57-59^. These analyses revealed relatively high level of PCYT1A and significantly low level of PCYT1B in monocytic AML (FAB M4 and M5, p value, <0.0001, 0.0002, respectively) compared to other AML subtypes (Fig. 5e, f). Taken together, we concluded that PCYT1A is a critical regulator in monocytic AML cells including those with MLL-rearrangements mainly due to the low level of PCYT1B in them.

**Fig. 5.**
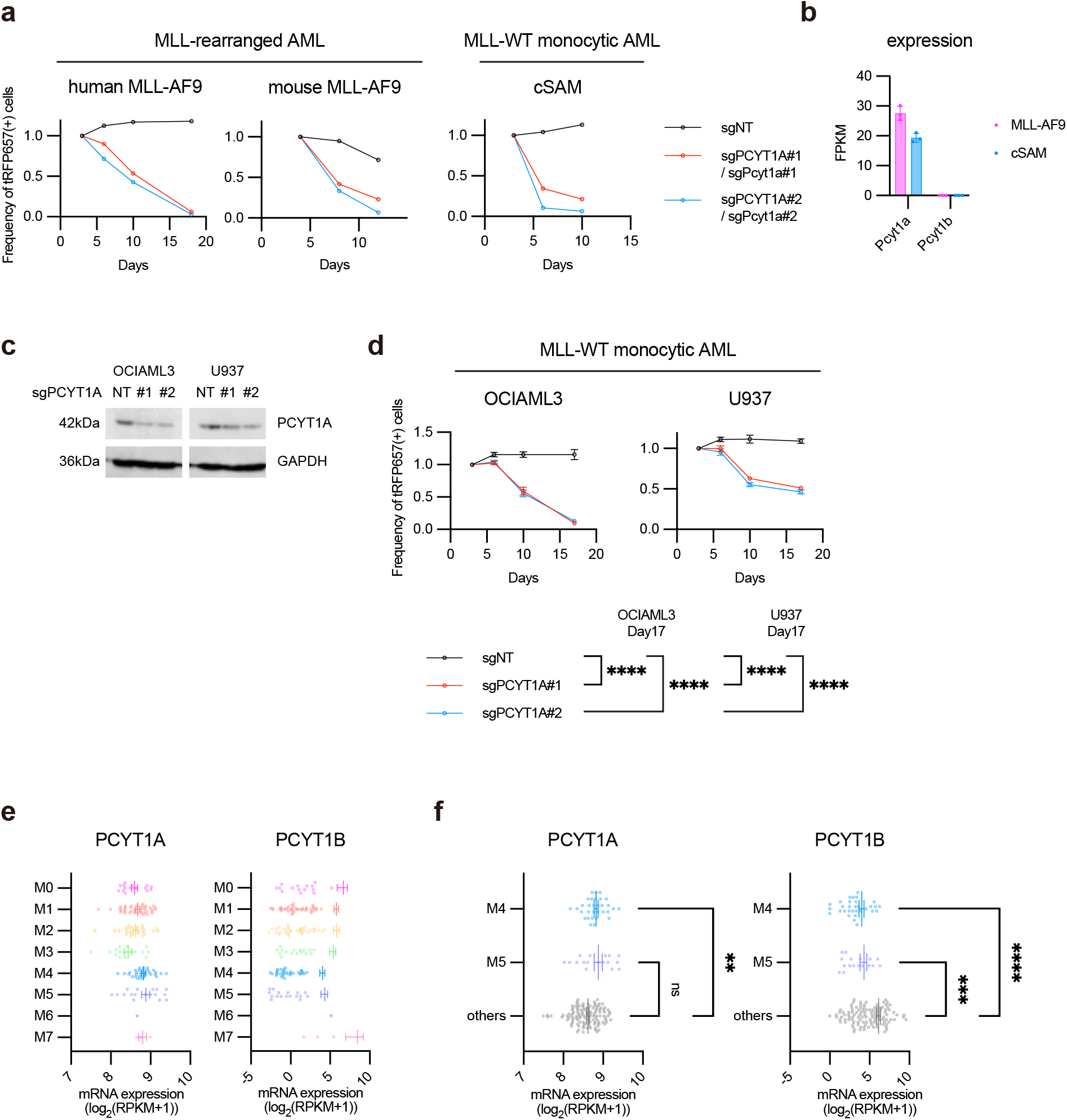
PCYT1A is essential for the growth of monocytic AML. (**a**) Human MLL-AF9 cells, mouse MLL-AF9 cells, and cSAM cells were transduced with Cas9 together with non-targeting (NT) sgRNA or PCYT1A/Pcyt1a-targeting sgRNAs (sgPCYT1A/Pcyt1a) co-expressing tRFP657. Changes in the frequency of tRFP657^+^ cells (sgRNA-transduced cells) in cell cultures are shown. Results are normalized to the frequency of tRFP657^+^ cells at day 4, set to 1. These MLL-AF9 expressing AML cells and MLL-WT monocytic AML cells are not registered in DepMap dataset. (**b**) FPKM values obtained via RNA-seq analysis. The data were retrieved from GSE135008 (MLL-AF9: n=3, cSAM: n=3)^67^. (**c, d**) OCIAML3 and U937 cells were transduced with Cas9 together with non-targeting (NT) sgRNA or PCYT1A-targeting sgRNAs (sgPCYT1A) co-expressing tRFP657. PCYT1A depletion by each sgRNA was confirmed. (**c**) Changes in the frequency of tRFP657^+^ cells (sgRNA-transduced cells) in cell cultures (triplicates) are shown. (**d**) Results are normalized to the frequency of tRFP657^+^ cells at day 4, set to 1. Data are shown as mean ± s.d. ns not significant, \**P*<0.05, \*\**P*<0.01, \*\*\**P*<0.001, \*\*\*\**P*<0.0001, two-way ANOVA multiple comparisons. (**e-f**) mRNA expression levels of PCYT1A and PCYT1B in 162 AML patients’ samples. The data were retrieved from cBioPortal and TCGA. M0 – M7 indicates the FAB classification. Data are shown as mean ± s.e.m. \*\**P*<0.01, \*\*\**P*<0.001, \*\*\*\**P*<0.0001, Brown-Forsythe and Welch ANOVA tests.

### Exploring therapeutic vulnerabilities in *TP53*-mutated AML

We then used our analytic system to identify novel therapeutic targets in *TP53*-mutated AML by comparing the Gene Effect Scores in eighteen *TP53*-mutated and eight *TP53*-WT AML cell lines registered in DepMap Public 22Q2. BCL2L1 (also known as BCL-XL) was ranked first (gap of Gene Effect Score, -0.676; unpaired t test P value, 0.0005), indicating that BCL2L1 is an important regulator specifically in *TP53*-mutated AML (Fig. 6a, b). To confirm this finding, we then assessed the effect of BCL2L1 depletion in *TP53*-mutated or *TP53*-WT AML cell lines. We transduced Cas9 together with NT or BCL2L1-targeting sgRNAs (coexpressing tRFP657) into two *TP53*-mutated AML cell lines (HEL and TF1) and two *TP53*-WT AML cell lines (MOLM13 and OCIAML3), and tracked the frequency of tRFP657-positive cells in culture. Efficient depletion of BCL2L1 was confirmed by western blotting (Fig. 6c). BCL2L1 depletion inhibited the growth of *TP53*-mutated cells, while it showed only a marginal effect on *TP53*-WT cells (Fig. 6d). These data suggest that *TP53*-mutated AMLs are highly dependent on BCL2L1.

**Fig. 6.**
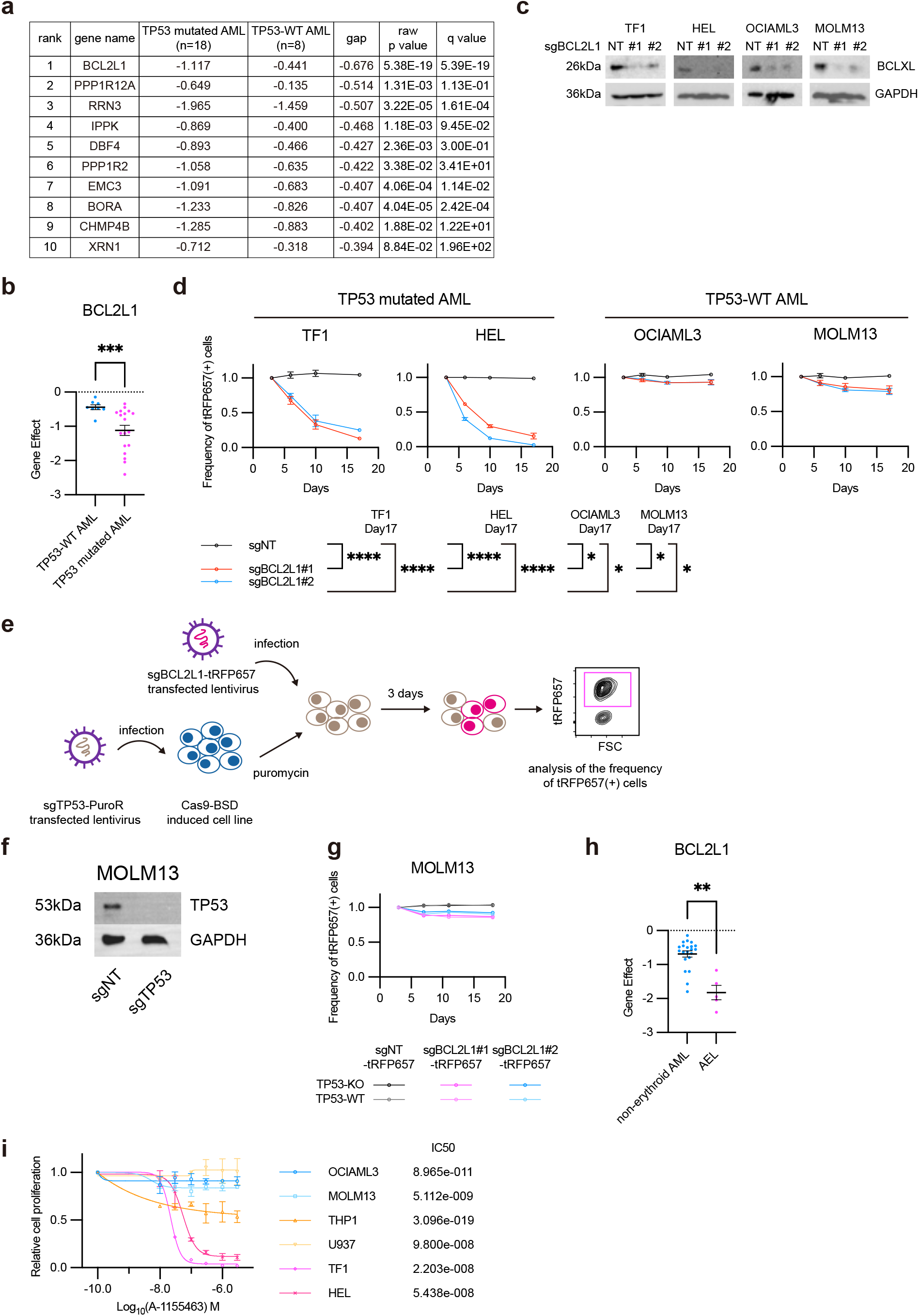
BCL2L1 is important for the growth of acute erythroid leukemia. (**a**) Results of DepMap Gene Effect Score analysis between *TP53*-mutated AML and *TP53*-WT AML. (*TP53*-mutated AML: n=8, *TP53*-WT AML: n=18) The top10 genes that are specifically essential in *TP53*-mutated AML are shown in the table. Unpaired t test with Welch’s correction and Benjamini-Hochberg procedure. (**b**) Dot plots of Gene Effect Scores for AML. Data are shown as mean ± s.e.m. \*\*\**P*<0.001 (**c-d**) *TP53*-mutated (TF1 and HEL) and *TP53*-WT (OCIAML3 and MOLM13) AML cells were transduced with Cas9 together with non-targeting (NT) sgRNA or PCYT1A-targeting sgRNAs (sgPCYT1A) co-expressing tRFP657. PCYT1A depletion by each sgRNA was confirmed in the AML cells (**c**). Changes in the frequency of tRFP657^+^ cells (sgRNA-transduced cells) in cell cultures (triplicates) are shown (**d**). Results are normalized to the frequency of tRFP657^+^ cells at day 4, set to 1 (**d**). Data are shown as mean ± s.d. \**P*<0.05, \*\*\*\**P*<0.0001, two-way ANOVA multiple comparisons. (**e**) Schematic presentation of experimental procedures for experiments shown in Fig. 6g. MOLM13 cells (coexpressing BSD) were transduced with non-targeting (NT) sgRNA or a sgRNA targeting TP53 [coexpreessing Puromycin Resistance (PuroR)]. After puromycin (1μg /mL) and blasticidin (10 μg/mL) selection, the cells were then transduced with NT or sgRNAs targeting BCL2L1 (coexpressing tRFP657). The frequency of tRFP657-positive cells was monitored by FACS starting 72 hours after the transduction of BCL2L1-targeting sgRNA. (**f**) TP53 depletion by each sgRNA was confirmed in MOLM13 cells. (**g**) Changes in the frequency of tRFP657^+^ cells (NT or sgBCL2L1-transduced cells) in cell cultures (triplicates) are shown. Note that BCL2L1-targeting sgRNAs did not inhibit the growth even in TP53-depleted MOLM13 cells. (**h**) Dot plots of Gene Effect Scores for AML. (AEL: n=5, non-erythroid AML: n=21) Data are shown as mean ± s.e.m. ns: not significant, \*\**P*<0.01. (**i**) Cell Viability Assays were performed using Cell Counting Kit-8. Two *TP53*-WT AML cell lines (OCIAML3 and MOLM13), two *TP53*-mutated AML cell lines with erythroid differentiation (TF1 and HEL) and two *TP53*-WT AML cell lines (THP1, U937) were treated with A-1155463 at the indicated concentration for 72h in duplicates. Data are normalized to vehicle control and are shown as mean ± s.e.m.

### BCL2L1 is a critical regulator in AEL

We next investigated whether TP53 is directly associated with the dependency on BCL2L1 in AML. To mimic the mutation-induced TP53 loss of function, we transduced Cas9 and a *TP53*-targeting sgRNA (sgTP53, coexpressing puromycin resistance gene) into a *TP53*-WT AML cell line MOLM13^28, 60^. The TP53-depleted MOLM13 cells were then used in the experiments to assess the effect of BCL2L1 depletion (Fig. 6e, f). Depletion of BCL2L1 showed no effect even in the *TP53*-depleted MOLM13 cells (Fig. 6f, g), suggesting that the increased susceptibility of *TP53*-mutated AML cells to BCL2L1 depletion is not the direct consequence of TP53 inactivation.

Because *TP53* mutations are common in AEL^24, 25^, we speculated that TF1 and HEL are sensitive to BCL2L1 depletion not because they have *TP53* mutations but because they have erythroid characteristics. To examine this possibility, we classified AML cells registered in DepMap Public 22Q2 into AEL and others. Interestingly, we found a more substantial difference in the Gene Effect Score of BCL2L1 (gap of Gene Effect Score, -1.132; unpaired t test P value, 0.0033) between AEL and other AMLs than that between WT and *TP53*-mutated AMLs (Fig. 6h). We then examined the effect of BCL2L1-selective inhibitor (A-1155463^35^) on the growth of two *TP53*-mutated AEL cell lines (TF1, HEL), two *TP53* mutated non-AEL cell lines (THP1, U937) and two *TP53*-WT AML cell lines (OCIAML3, MOLM13). As expected, the *TP53*-mutated AEL cell lines (TF1 and HEL) were susceptible to the BCL2L1 inhibitor, while two *TP53*-WT AML cell lines (OCIAML3, MOLM13) were not. Importantly, the two non-AEL cell lines (THP1 and U937) were not sensitive to the BCL2L1 inhibitor despite the presence of *TP53* mutations in them (Fig. 6i). These results suggest that the erythroid differentiation, but not the presence of *TP53* mutations, confers dependency to BCL2L1 in AML.

## DISCUSSION

Since the public release of the DepMap, numerous studies have been conducted to identify genes required for survival and proliferation in specific cancer subtypes^9-12^. Although DepMap provides several cell line groups, they are not sufficient to explore therapeutic targets in defined cancer subtypes with specific mutations or fusions. In this study, we developed a two-group comparison system to identify key genes that are essential in a specific group of cancer cell line with a particular genetic abnormality, which would contribute to identify therapeutic vulnerabilities in each cancer subtype.

Using this system, we explored specific vulnerabilities in two AML subtypes with poor prognosis. The analyses led to the identification of PCYT1A and BCL2L1 as a critical regulator in MLL-rearranged AML and *TP53*-mutated AML, respectively. Interestingly, further investigation revealed that PCYT1A is not a selective regulator in MLL-rearranged AML, but is important for the growth of monocytic AMLs “including” those with MLL-rearrangements. Similarly, BCL2L1 is essential in AMLs with erythroid differentiation in which *TP53* is frequently mutated. These results strongly suggest the importance of cell of origin, rather than the genetic aberrations alone, to identify subtype-specific therapeutic targets in AML. Although genetic information is being gradually incorporated into classification and treatment decisions for AML, our study reinforces the need to retain the FAB classification to provide optimal therapies for each AML subtype.

PCYT1A is a rate-limiting enzyme to synthesize the phosphatidylcholine during lipid metabolism. PCYT1A is often upregulated through RAS and MYC pathways in various types of cancer, and is thought to promote cancer cell growth^61-63^. Although PCYT1B has similar enzymatic activity to PCYT1A, the role of PCYT1B in cancer is not well understood. Interestingly, we found that AMLs with monocytic features including MLL-rearranged AML were particularly sensitive to PCYT1A depletion due to the low PCYT1B expression in them. In contrast, non-monocytic AML cells grew normally after PCYT1A depletion because they express sufficient PCYT1B to compensate for the loss of PCYT1A. Thus, our findings suggest that the “low” expression of PCYT1B could be a biomarker to predict the susceptibility of each cancer to PCYT1A inhibition. Such synthetic lethal strategy targeting metabolic pathways could be a promising approach to treat cancers that are overly dependent on a particular metabolic gene.

BCL2L1 is a member of the anti-apoptotic BCL2 family proteins, such as BCL2, BCL2L1 and MCL1^64^. The upregulation of the anti-apoptotic BCL2 family members is frequently observed in various types of cancer. Many hematopoietic neoplasms rely on BCL2 to evade apoptosis and are sensitive to BCL2-selective inhibitor venetoclax^65^. However, recent clinical studies have shown that *TP53*-mutated AMLs including AEL are resistant to the venetoclax-based therapies. In this study, we found that AEL cells including those with *TP53* mutations are highly dependent on BCL2L1 for their growth. While preparing this paper, another group also reported the critical role of BCL2L1 in AEL and acute megakaryoblastic leukemia^66^. Our data confirmed their findings, providing a rationale for the clinical development of BCL2L1-specific inhibitors to treat AEL.

In summary, we developed a two-group comparison system to identify therapeutic vulnerabilities in specific cancer subtypes using the information provided by DepMap. We applied this system to AML and identified PCYT1A and BCL2L1 as a promising target to treat monocytic AML and AEL, respectively. This method could accelerate the discovery of subtype-specific therapeutic targets in diverse cancers in future studies.

## ACKNOWLEDGEMENT

We thank C. Kamiya, R. Shimura for their excellent technical assistance, and Kitagawa lab members for constructive feedback on the DepMap analysis system. We are also grateful to the FACS Core at the Institute of Medical Science, The University of Tokyo. This work was supported by Grant-in-Aid for Scientific Research (B) (22H03100, SG), Grant-in-Aid for Challenging Research (Exploratory) (22K19540, SG), Fund for the Promotion of Joint International Research (Fostering Joint International Research (B)) (22KK0127, SG), AMED under Grant Number (21ck0106644h0001, SG), Grant-in-Aid for Early-Career Scientists (JP22K16319, KY).

